# Hidden diversity and expanded host range of sarthroviruses, including terrestrial vertebrates

**DOI:** 10.64898/2026.06.16.732546

**Authors:** Ethan Mandojana, Lauren Lim, Julien Mélade, Josephine Rieken, Jane Hall, Mary Petrone, Jonathon C.O. Mifsud, Ezequiel M. Marzinelli, Karrie Rose, Edward C. Holmes, Kate Van Brussel

## Abstract

The *Sarthroviridae* are a family of highly compact satellite RNA viruses comprising one recognised species, extra small virus (XSV). *Macrobrachium rosenbergii* nodavirus (MrNV) is the associated helper virus of XSV and their co-infection has been linked to white tail disease in freshwater prawns globally, although the role of XSV is remains unclear. Here, we describe the discovery and characterisation of ten novel, highly divergent sarthrovirus species from a range of hosts and environments within a small geographical region in Australia. These comprise novel sarthroviruses associated with marine sponges, seal and dingo faeces, environmental marine sediment samples and Indo-Pacific geckos (*Hemidactylus garnotii*). All the novel viruses possess only a capsid protein, consistent with the genome of XSV, yet exhibit substantial sequence divergence. Notably, some sarthrovirus variants seem to utilise different replication systems despite being genetically identical and present in the same host species. Sequences from nodaviruses, which could plausibly act as helpers, were associated with some, but not all, the sarthroviruses identified here. Phylogenetic analyses support the expansion of the *Sarthroviridae* into multiple distinct lineages, comprising at least seven genera. Collectively, these findings reveal a broader ecological distribution and evolutionary diversity of sarthroviruses and highlight the possibility of alternative replication strategies and tissue tropism in diverse animal host.

**Significance:** Sarthoviruses are small (∼800 nucleotides) satellite RNA viruses associated with a nodavirus of crustaceans that acts as a helper. To date, the only known sarthovirus is extra small virus (XSV), which also represents the sole species within the *Sarthroviridae*. Here, we report the detection of ten divergent sarthroviruses sampled from diverse animal hosts, including vertebrates, that expand the family to 11 species and at least seven genera. These viruses were detected from various host taxa and environmental samples from a confined geographical region in eastern Australia, suggesting that they are ecologically connected. Notably, we did not detect nodaviruses in all samples containing sarthroviruses, suggesting that different viruses may act as helpers for sarthovirus replication.

## Introduction

Satellite RNA viruses lack essential genes such that they depend on the co-infection of a larger helper virus for replication, assembly and/or cell entry. Satellite RNA viruses are characterised by single-stranded, small and simple genomes that are genetically distinct from their associated helper viruses, and have arisen multiple times independently. Arguably the best known helper-satellite RNA virus pair in vertebrates belongs to members of the *Kolmioviridae* (i.e., the ‘delta-like’ viruses). In humans, hepatitis D virus is intimately associated with co-infection with hepatitis B virus, although other vertebrate kolmioviruses seemingly utilize a variety of viruses as helpers (1–4). Hence, it is unknown whether satellite-helper relationships and replication strategies remain stable through evolutionary time.

Sarthroviruses (*Sarthroviridae)* possess a highly compact RNA genome of approximately 800 nucleotides that encodes only a capsid protein and lacks the replicative machinery typical of autonomous RNA viruses, categorising them as satellite viruses (5). The sole sarthrovirus characterised to date – extra small virus (XSV) – was first identified by transmission electron microscopy in giant freshwater prawns (*Macrobrachium rosenbergii*) during an investigation of white tail disease (6). The virus was observed next to a larger virus particle that formed crystalline structures, now known to be *Macrobrachium rosenbergii* nodavirus (MrNV) (7–9). The consistent co-detection of XSV with MrNV across multiple affected individuals established MrNV as the helper virus of XSV, and their co-infection has been proposed as the cause of the disease (6–9). Despite being recovered in crustacean hosts from multiple countries, all XSV sequences share 98-100% amino acid identity over the capsid protein and constitute a single recognised species (5). Although the *Sarthroviridae* are currently placed within the order *Tombendovirales* alongside satellite RNA viruses of plants, there is no detectable sequence homology between the XSV capsid protein and those of other viruses. To date, sarthroviruses have not been naturally associated with hosts outside of crustaceans and aquatic insects, such that their evolutionary origins and host range remain largely unknown.

A major barrier to understanding the diversity and evolution of sarthroviruses has been the limited scope of host taxa surveyed. Herein, we describe the discovery and characterisation of ten highly divergent sarthroviruses from New South Wales, Australia, sampled in phylogenetically diverse hosts. This includes one found in the liver tissue of a terrestrial vertebrate host, the Indo-Pacific gecko (*Hemidactylus garnotii*). Additional sarthroviruses, including relatives to those found in Indo-Pacific geckos, were recovered from environmental and faecal samples collected within the same geographic region, revealing a previously unrecognised breadth of host and ecological associations for this virus family. Together with the co-detection of nodaviruses and other RNA viruses, these results expand the known diversity, host range, and evolutionary context of sarthroviruses.

## Results

### Expansion of the *Sarthroviridae* to include seven genera

Five geckos, one seal faecal sample and four sea sponges underwent metatranscriptomic sequencing for the detection of sarthroviruses (Table S1). In addition, we mined the NCBI SRA database and identified sarthroviruses in 21 SRA libraries (19 marine sediment, one dingo faeces and one common carp) that were also included in the analysis for this study (Table S1). Overall, the metagenomic sequencing performed as part of this study combined with the SRA mining yielded ten novel sarthroviruses in various hosts (see below). To highlight their small size, and prior to formal classification by the International Committee on Taxonomy of Viruses (ICTV), they were tentatively named gimli virus 1 and gimli virus 2, half pint virus, incy wincy virus, itsy bitsy virus, pee wee virus, teeny tiny virus, teeny weeny virus, titchy virus and very small virus. All these novel viruses contained only a nucleocapsid CP confirmed by VirSorter2 and geNomad, consistent with the genomic structure of the previously identified XSV.

Strikingly, itsy bitsy virus was detected in 17 SRA data sets, two gecko liver and three sea sponge sequencing libraries, all with 96 - 100% nucleotide similarity, while two SRA libraries contained half pint virus and three contained pee wee virus. In the seal faecal sequencing library two variants of teeny weeny virus were identified that share 86.6% nucleotide and 91.6% amino acid identity. Similarly, two species of gimli virus were identified in two SRA libraries that share 85.6% nucleotide and 85.9% amino acid identity. Comparison involving all ten novel sarthroviruses revealed amino acid identities ranging from 14% to 66%, and hence indicative of substantial diversity (Figure 1). Phylogenetic analysis of the translated amino acid sequences of the capsid protein, combined with amino acid similarities, suggested that the *Sarthroviridae* contains at least 11 species and seven genera, although the evolutionary relationships among the genera were often uncertain due to low bootstrap values (Figure 1). Maximum likelihood trees were also generated using trimmed alignments. As the species groups remained the same with minor repositioning of the clades, and with low internal nodes bootstrap support, they are not shown here. While the capsid amino acid alignment was complicated and did not highlight substantial sequence homology, HHpred matched all capsid amino acid sequences to XSV with a probability of between 93-100%, E-values ranging from 0.19 to 2.8e-76 and coverage of 45 to 98%. BLASTN and BLASTX searches using the nucleotide sequence of the capsid protein yielded no hits in the nt and only hits to XSV capsid protein in the nr databases, suggesting the source of the novel sarthroviruses is not host related. Attempts to culture itsy bitsy virus and its helper in mosquito (C6/36), canine (MDCK), fish (CHSE-214) and tick (HAE/CTVM8) cell lines were unsuccessful (results not shown) and in the absence of this data we were unable to determine whether itsy bitsy virus is capable of forming virions.

**Figure 1.**
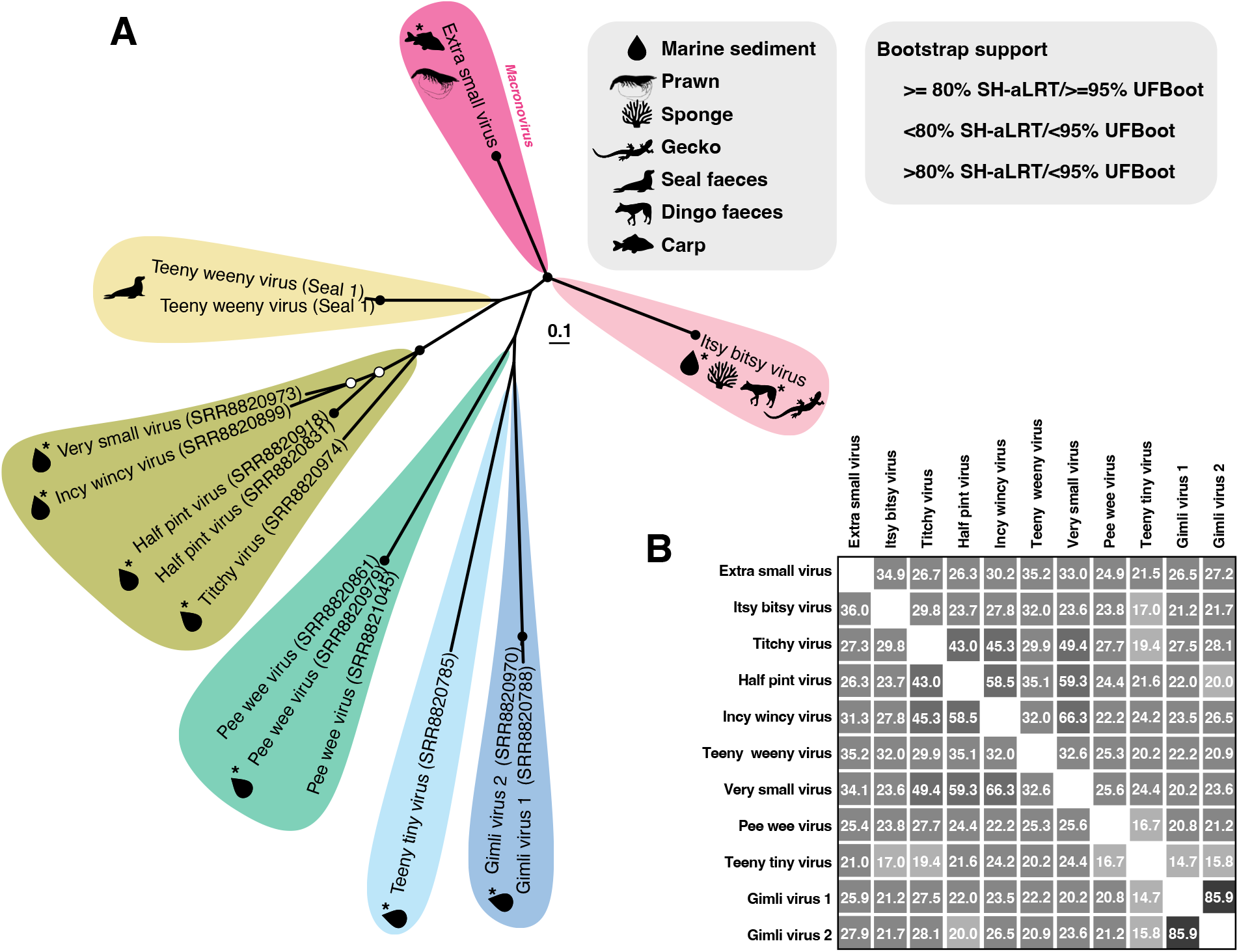
Phylogenetic relationships and genetic distances among the *Sarthroviridae*. (A) Unrooted maximum likelihood phylogenetic tree of the complete capsid protein MAFFT alignment of all 14 available NCBI GenBank XSV sequences and the novel species identified in this study. The XSV and itsy bitsy virus clades have been collapsed in the tree for better visualisation. Each virus sequence has the sequencing library in parentheses and the asterisk represents the sequences identified in libraries taken from the NCBI SRA database. All horizontal branch lengths are drawn to a scale of amino acid substitutions per site and the support values (SH-aLRT and UFBoot) are shown at the nodes. (B) Amino acid genetic distances of all the novel sarthroviruses identified in this study and XSV using MAFFT.

The marine sediment, gecko, sea sponge and seal faecal samples were all collected from New South Wales (NSW) Australia, with the marine sediment, sea sponge and gecko samples all collected from suburbs in Sydney during 2016 - 2026 (Figure 2). The seal and dingo (SRR32313011) were the only individuals that were not located in the Sydney area (Figure 2). An additional five Indo-pacific geckos from Sydney had their liver RNA screened for itsy bitsy virus with only one other gecko being PCR positive. Based on the ten geckos screened for this study the approximate frequency of itsy bitsy virus in the Sydney area is 30% (3/10).

**Figure 2.**
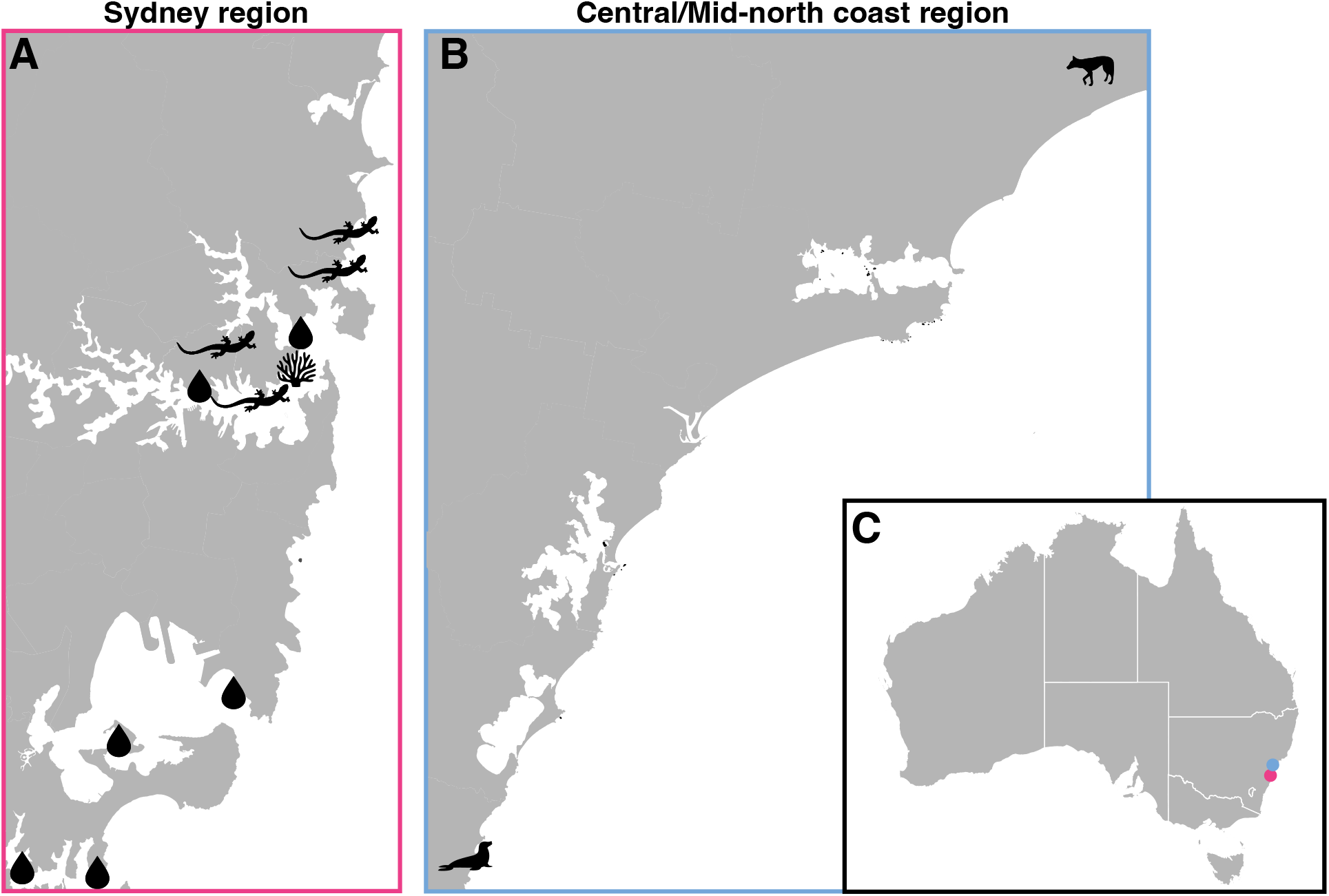
Map of the sampling locations in NSW, Australia. (A) Sampling locations for each animal host for the Sydney region and (B) sampling locations in the Central/Mid-North coast region of NSW. (C) Map of Australia showing the corresponding Sydney and Central/Mid-North coast sampling regions depicted by a circle. An image for each sampled host is attached to each location.

### Multiple potential virus-helper relationships in the *Sarthroviridae*

As XSV is replication incompetent without co-infection with a nodavirus, we next screened all metatranscriptomic libraries for *Nodaviridae* contigs. Accordingly, nodavirus sequences were detected in one of two geckos, the seal faeces, and all three sea sponge libraries. While no nodavirus sequences were detected in one of the gecko libraries, four nodavirus reads of 139 bp were detected by Kraken2, suggesting that a nodavirus was most likely present but was missed during RNA sequencing due to low total RNA yield. The additional gecko that did not undergo metatranscriptomic sequencing but was PCR positive for itsy bitsy virus similarly did not contain the same gecko nodavirus. Unsurprisingly, the NCBI SRA marine sediment samples contained >400 nodavirus contigs, such that only contigs greater than 600bp were assessed.

Interestingly, the two Indo-pacific gecko nodaviruses do not cluster together in the RdRp phylogeny but instead fall within different lineages (Figure 3). However, as the sequence of Indo-pacific gecko nodavirus 1 is only 46 amino acids in length its phylogenetic positioning cannot be accurately determined. The two nodaviruses in the seal faecal library were identified as nodaviruses of fish, suggesting that the source of these viruses and potentially teeny weeny virus is likely dietary (Figure 3). The nodaviruses detected in the SRA libraries were positioned throughout the two lineages (Figure 3). Finally, we did not detect MrNV in the NCBI SRA library (SRR13709132) that contained XSV, again likely due to sequencing depth (10).

**Figure 3.**
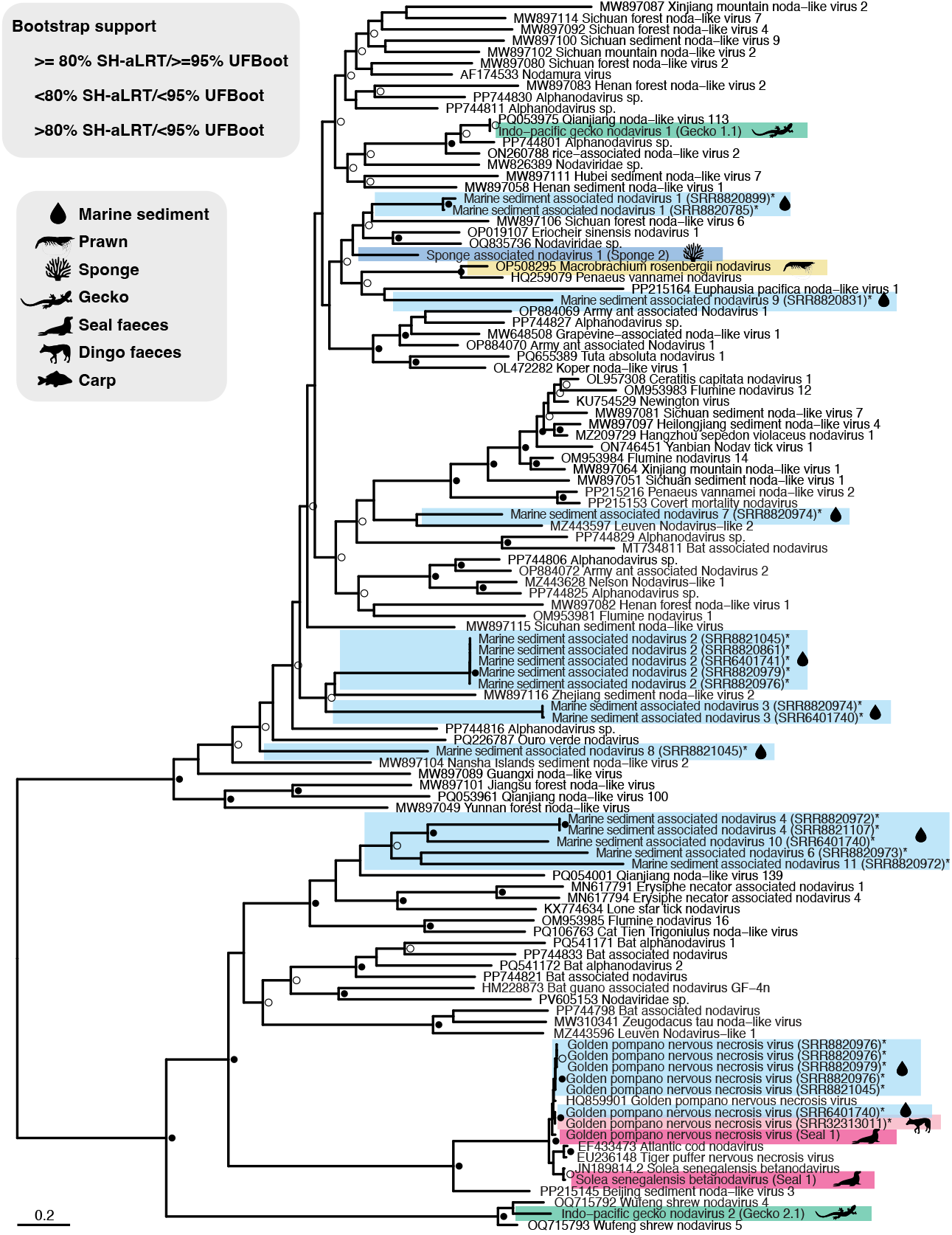
Phylogenetic relationship of the *Nodaviridae*. Maximum likelihood phylogenetic tree of the RdRp MAFFT alignment of available NCBI GenBank nodavirus sequences and the sequences identified in this study. Each virus sequence has the sequencing library in parentheses and the asterisk represents the sequences identified in libraries taken from the NCBI SRA database. Unaligned sections and gaps were removed from the alignments before tree construction. All horizontal branch lengths are drawn to a scale of substitutions per site and the support values (SH-aLRT and UFBoot) are shown at the nodes. The tree is midpoint-rooted for clarity.

As well as the nodaviruses, we assessed whether the gecko libraries harboured other viruses. Accordingly, astrovirus contigs were detected in one gecko lung library but were only 300bp (RdRp) and 401bp (capsid) in length (Figure S1). In addition, dicistrovirus and paramyxovirus contigs were identified in one gecko library, although subsequent PCR revealed that the dicistrovirus contigs were present in the gecko not infected with itsy bitsy virus and that the paramyxovirus was an endogenous virus element (Figure S2). Kraken2 analysis identified reads mapping to the genus *Raillietiella*, a parasitic crustacean that is known to infect the lungs of species in the *Hemidactylus* genus (11, 12). *Raillietiella* reads were found in all four gecko libraries that underwent RNA-Seq, including both the lung and liver libraries, with read counts ranging from 30,722 to 20,5085.

### Sarthrovirus capsid protein structures

We used AlphaFold 3 to predict the protein structures for all ten novel species. All novel viruses, excluding gimli virus 2, showed high average pLDDT values (73.36% to 91.81%) and formed a single jelly roll fold also seen in the experimental protein structure of XSV (PDB 6JJA) as well as other plant satellite viruses (Figure 4). The predicted template modelling (pTM) score for all sequences except gimli virus 2 were above 0.5, suggesting the predicted structure is likely reflective of the true structure. Foldseek assigned the highest probability and lowest E-value hit in the PDB100 database as XSV (PDB 6JJA) for all sequences again with the exception of gimli virus 2. Interestingly, AIUPred identified two binding regions in titchy virus at the beginning and end of the amino acid sequence, although only the beginning of the sequence was identified as a highly disordered region. This 3’ end of the titchy virus capsid protein is likely the C-terminal region and may potentially play a key role in replication and helper virus interaction. Overall, the protein structure analyses suggest that nine of the ten novel sequences presented in this study share structural homology with XSV.

**Figure 4.**
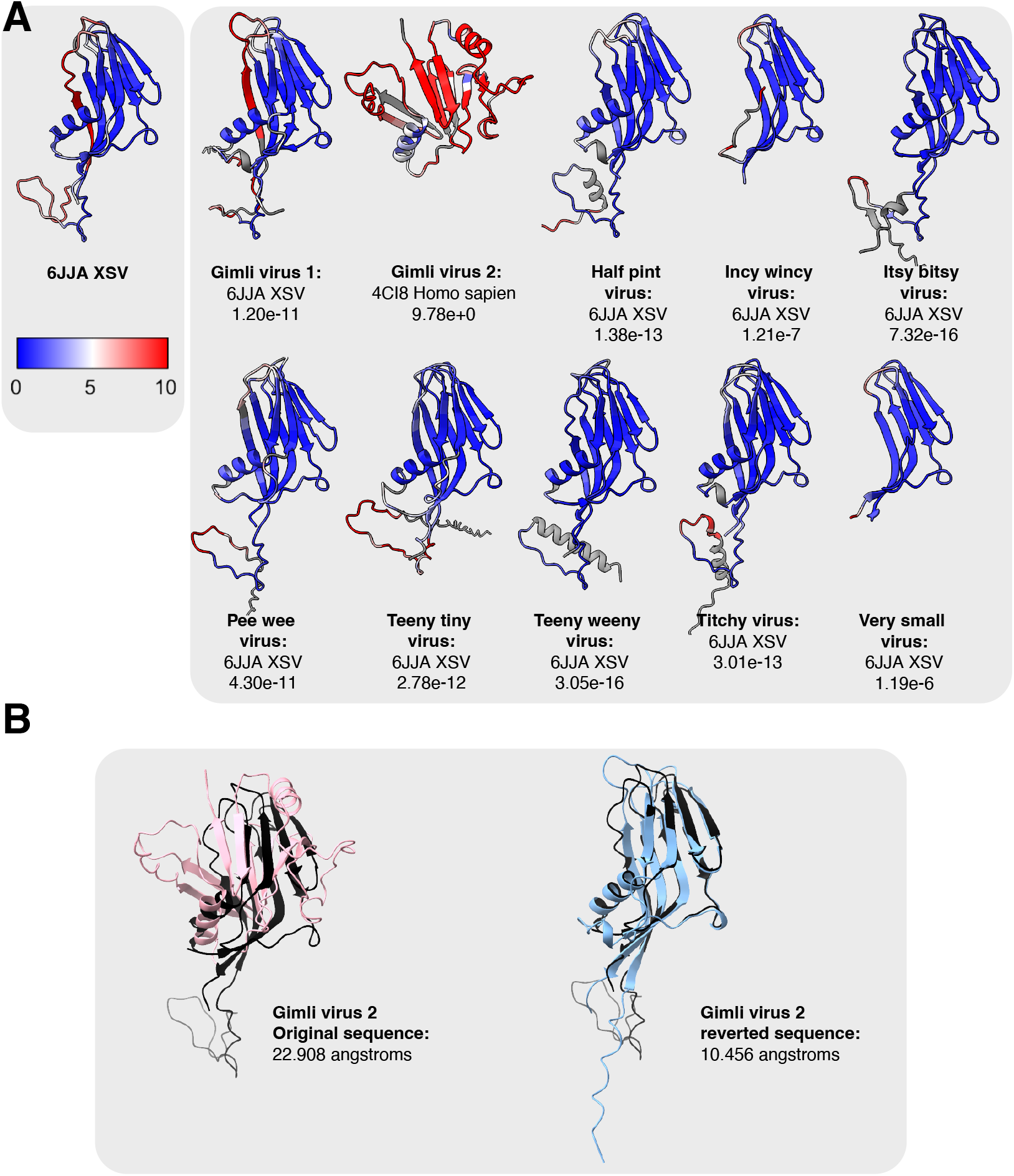
Predicted protein structure of novel Sarthroviruses. (A) Protein structure prediction of sarthroviruses obtained by AlphaFold 3 and coloured by the root-mean square deviation (RMSD) distance from 6JJA XSV. (B) Gimli virus 2 predicted protein structure superimposted over 6JJA XSV (coloured in black).

The low average pLDDT value (35.97%) and pTM score (0.21) for gimli virus 2 meant that we were unable to assign confidence to the protein structure generated by AlphaFold 3. Despite the lack of an accurate predicted protein structure, as gimli virus 1 and gimli virus 2 share 86% amino acid identity over their capsid gene and HHpred matched a section of its sequence to XSV (PDB 6JJA), we believe that gimli virus 2 is a sarthrovirus. We therefore investigated why AlphaFold 3 was able to confidently predict a protein structure for gimli virus 1 but not gimli virus 2. An obvious different between the two is the 32 amino acid sequence at the beginning of gimli virus 1, likely the N-terminal region of the protein and which was predicted to likely be a disordered binding region by AIUPred. A highly disordered or binding region was not identified in the amino acid sequence of gimli virus 2. Additionally, we used both the hydrophobicity and amino acid alignments to locate three amino acid substitutions in gimli virus 2 that resulted in a hydrophobic/hydrophilic/neutral amino acid change compared to gimli virus 1. When all three substitutions in gimli virus 2 were reverted to the residues present in gimli virus 1 the average pLDDT confidence was 82.26%. The predicted protein structure with the reverted amino acid sequence aligned completely in the *α* helix and *β* sheet with XSV (PDB 6JJA, Figure 4). Foldseek was also able to assign the PDB100 database hit as XSV (PDB 6JJA), with an E-value and sequence id comparable to gimli virus 1. These amino acid positions in gimli virus 2 are potentially located in regions of the sequence that provide structure to the protein and may play a role in protein stability in nature.

## Discussion

Herein we demonstrate the occurrence of a sarthrovirus in a terrestrial vertebrate host, as well as those marine animals and environmental samples, thereby expanding the known host range in this unusual group of satellite RNA viruses beyond crustaceans and insects. In addition, the discovery of ten highly divergent sarthroviruses across host taxa and within a geographically confined region of NSW Australia indicates that these viruses occupy a broader ecological niche than previously recognised.

Protein structure predictions were able to confidently assign nine of the ten novel viruses as sarthrovirus-like, all possessing a jelly roll fold consistent with the experimental structure of XSV (PDB 6JJA). Gimli virus 2 was the only sarthrovirus that did not produce a confident structural prediction using AlphaFold 3 despite its high sequence similarity to gimli virus 1 (85%). The truncated sequence of gimli virus 2 and inability of AIUPred to locate a highly disordered region suggests that the gimli virus 2 sequence is missing the N-terminal region of the capsid protein. In addition, the sequences assembled for very small virus and incy wincy virus were truncated at the beginning of the capsid protein and are ∼60 aa shorter than gimli virus 2, although AlphaFold 3 was able to produce confident structures for both viruses. The properties of the N-terminal region of these sarthroviruses and the potentially C-terminal region in titchy virus identified by AIUPred require further investigation to determine their influence on replication and helper virus interaction.

Although XSV is currently the only classified member of the *Sarthroviridae*, this study has substantially expanded the diversity of this family, with phylogenetic analysis of the capsid and measurements of genetic diversity indicating that the *Sarthroviridae* contains at least 11 species that can be separated into seven genera, although the evolutionary relationships among them are harder to resolve. Indeed, itsy bitsy virus shares only 37% amino acid identity with known XSVs, while teeny tiny virus, gimli virus 1 and 2 and pee wee virus share only ∼20% to all other sarthroviruses, suggesting that these reflect ancient divergence events (with the caveat that nothing is known about the rate of evolutionary change in these viruses) and that the full host range of this virus family has yet to be documented. Interestingly, very closely related itsy bitsy virus sequences (96-100% similarity) were detected in samples from chordate, porifera and environmental sources, all sampled within close proximity from different years (2016 – 2026). Hence, it is possible that sarthroviruses undergo slower evolution than other RNA viruses, potentially attributable to their dependence on the presence of another virus for replication.

The close association between XSV and MrNV has been shown in studies involving cell culture experiments and electron microscopy, comprising one of the few documented examples of an RNA virus requiring a helper virus for replication (5, 8, 13), and observed over a large geographical range (6–9). Hence, we would expect to see that samples with identical sarthroviruses should also share nodaviruses, yet this was not the case here. The identification of itsy bitsy virus and a nodavirus concurrently in the liver of two individual Indo-Pacific geckos suggests that the functional dependency of sarthroviruses on helper nodaviruses is maintained in vertebrate hosts, although this requires experimental verification. The liver RNA from a third gecko (that was not sequenced) was PCR positive for itsy bitsy virus and negative for Indo-pacific gecko nodavirus 2. As a consequence, in the absence of experimental confirmation we cannot definitively conclude that nodaviruses are the helper viruses for our novel sarthroviruses. The presence of additional viruses in the gecko libraries, such as astroviruses and endogenised viruses, prevents definitive conclusions to be made on the possible helper virus(es) for itsy bitsy virus. Interestingly, the four satellite viruses of the *Tonesaviridae* are associated with different helper viruses of the same viral family, while satellite tobacco mosaic virus (*Tomosaviridae*) can utilise non-preferred helper viruses of the same viral genus but at the expense of replication efficiency (14, 15). Demonstration that all sarthroviruses require nodaviruses for replication would represent a remarkable conservation of a helper–satellite virus association across invertebrate and vertebrate host lineages. Conversely, demonstrating that these novel sarthrovirueses utilise helper viruses from different viral families would represent a case in which satellite RNA viruses share evolutionary origins but have evolved to use varstly different helper viruses. If this were the case it would potentially indicate that these novel sarthroviruses should be classified into different families within the same viral order.

The transmission source of XSV has not been fully identified, although it has been hypothesised that insects may act as vectors for virus spread to susceptible prawns (16). In particular, XSV and MrNV have both been detected in the aquatic insect *Belostoma sp, Aeshna sp* and *Notoncita sp* which can replicate together in mosquito cells (C6/36), raising the possibility of insect-mediated transmission between aquatic and terrestrial hosts (16, 17). However, the detection of sarthroviruses from marine sediment samples questions this hypothesis. It is plausible that insects or their larvae could be collected during sampling of marine sediment, although we did not observe high read counts for the Insecta class for this sample type (Figure S3). XSV transmission in susceptible prawns has been attributed to vertical transmission, while in brine shrimp it can occur through contact with an infected water source (5, 18). It is therefore plausible that transmission via a contaminated water source could be the cause of detection of itsy bitsy virus in geckos. Overall, however, insect transmission remains the most likely explanation for the detection of sarthroviruses in the terrestrial vertebrates although environmental transmission from water sources cannot be excluded. The detection of two fish nodaviruses and teeny weeny virus in the faecal sample of an Australian fur seal further suggests the source of this virus is fish, the primary dietary component of seals (19). XSV and MrNV have been previously detected in common carp samples from the Murray River, and an XSV with no MrNV was detected in a sequencing library from that same study, although this is hypothesised to represent environmental contamination (10).

Together, these findings reveal an unexpected diversity of sarthroviruses and substantially expand the known evolutionary and ecological breadth of the family *Sarthroviridae* and satellite RNA viruses. The repeated recovery of closely related sarthroviruses from diverse host and environmental samples collected within a confined geographic area in NSW, Australia over multiple years demonstrates their local persistence. The detection of sarthroviruses across different host taxa and environments could help answer key questions around virus interactions, ecology and evolution.

## Materials and Methods

### Sample collection and animal ethics

Samples were collected under ethics approval number 4c/10/21 provided by Taronga Conservation Society Australia’s Animal Ethics Committee and under the auspices of scientific license SL100104 issued by the New South Wales Department of Energy, the Environment, Climate Change and Water. Liver and lung tissue from ten Indo-Pacific geckos was collected from the Sydney region in NSW, Australia during April 2025 to March 2026 (Figure 2). A faecal sample was collected post-morten from an Australian fur seal (*Arctocephalus pusillus doriferus*) at Avoca Beach in the Central/Mid-North coast region of NSW in 2021 (Figure 2). Finally, sea sponge samples were collected from separate pier-pilings at ∼1.5-2m depths in Chowder Bay in Sydney Harbour during 2023 (Figure 2). The seal, all sea sponges and 5/10 geckos were submitted for metatranscriptomic sequencing at the Australian Genome Research Facility (AGRF).

### RNA processing, sequencing and read processing

Total RNA was extracted from gecko liver and lung tissue, seal faeces and sea sponges using the RNeasy Plus Mini Kit (Qiagen) following the manufacturer’s protocol. Total RNA was used to produce sequencing libraries with the Illumina Stranded Total RNA Prep with Ribo-Zero Plus rRNA depletion that were sequenced on the Illumina NovaSeqX platform (150bp paired-end) at the Australian Genome Research Facility (AGRF). The three sea sponge libraries were sequenced on the Illumina NovaSeq 6000 platform, S4 lane (150bp paired-end) at the AGRF. Raw read quality was assessed using FastQC and reads were quality filtered with sequencing adapters removed using Cutadapt (version 5.0) (20). Negative control libraries were run alongside the gecko, sea sponge and seal faecal libraries to confirm extraction kit reagents and columns were not the source of the identified viruses.

Filtered reads were *de novo* assembled into contigs using MEGAHIT (version 1.2.9) and these contigs were compared against the Reference Viral Database (RVDB) using DIAMOND BLASTX (version 2.1.6) (21–24). Any contigs that produced viral hits after RVDB screening were compared against the full NCBI non-redundant (nr) protein database using DIAMOND (version 2.1.6) (24). Viral genomes were extracted from these results and annotated in Geneious (version 2025.1.2) using a combination of Find ORFs, NCBI conserved domain database, VirSorter2, GeNomad and InterProScan (25–28). TaxonKit was used to assign taxonomic lineage information to the metadata output generated by database comparisons (29). To exclude environmental contamination of sequencing libraries, filtered reads were assigned to a taxonomic classification using Kraken2 (version 2.1.3) and virus read abundance was calculated using RSEM (version 1.3.3, Figure S3) (30, 31). A combination of HHpred, DeepVirFinder and geNomad were used to assess whether the novel sarthroviruses were indeed viral (25, 32, 33). Accordingly, sequences were considered to be of viral origin if they produced a confidence score of >0.7 and p-value of <0.5 for DeepVirFinder and/or a confidence of >0.7 for geNomad. Any sequence that produced a p-value >0.05 for DeepVirFinder or a confidence of <0.7 for geNomad was flagged as potentially viral.

### Sequence Read Archive (SRA) screening

To better understand the genetic diversity of the *Sarthroviridae*, XSV and the novel gecko and seal sarthroviruses were used to mine the SRA database for any metatranscriptomes with hits to these sarthroviruses using the PebbleScout Metagenomic Volume 1 and 2 databases (34). Only SRAs with a coverage above 5% were screened for sarthroviruses, resulting in the identification of 51 libraries (see Results). As PebbleScout does not currently contain SRAs submitted to the database after 2023, an additional 586 reptile liver and water libraries from 2024 to 2026 were downloaded and mined. Kingfisher was used to download the sequencing libraries from NCBI, which were then parsed through the same virus discovery pipeline as above (35).

### Protein structure prediction and phylogenetic analysis

AlphaFold 3 was used to predict protein structures for the 10 novel sarthroviruses identified here using the default settings and the resulting structures were compared to the PDB100 database using Foldseek (36, 37). The predicted protein structures generated by AlphaFold 3 were visualised in UCSF ChimeraX for publication (36, 38). Disordered, binding and linker regions in the amino acid sequences were predicted using AIUPred (39, 40).

Reference protein sequences for each viral family were downloaded from NCBI GenBank and aligned with the virus sequences from this study using the E-INS-I algorithm within MAFFT (version 7.490) in Geneious (version 2025.1.2) (41). Sequence alignments for the *Astroviridae, Dicistroviridae* and *Nodaviridae* were trimmed using trimAl (version 1.5), employing the gappyout flag to remove poorly aligned regions and gaps (42). IQ-TREE (version 2.3.6) employing the Model Finder Program was used to infer maximum likelihood trees using amino acid sequence alignments (43, 44). Nodal support was estimated using 1000 SH-like approximate Likelihood Ratio Test (SH-aLRT) and UltraFast Bootstrap (UFBoot) replicates (45). Nearest neighbour interchange (NNI) was used to control for overestimation of UFBoot values.

### PCR screening sarthroviruses and nodaviruses

Total RNA was used to perform reverse transcription PCR using the SuperScript IV One-Step RT-PCR following the manufacturer’s protocol. Primers were designed and sanger sequencing output trimmed and aligned using Geneious Prime (Table S2). Amplicon was sequenced at the AGRF.

## Data Availability

The raw sequence reads generated in this study are available at the NCBI SRA under BioProject XXX accessions XXX-YYY. Virus consensus sequences have been deposited in the NCBI GenBank database under accession numbers XXX-YYY.

## Acknowledgements

This work was funded by a National Health & Medical Research Council (Australia) Investigator grant (GNT2017197) and an Australian Research Council Discovery Project grant (DP240101313) to ECH, and a Princeton University Streicker International Fellowship to EM. The authors acknowledge the technical assistance provided by the Sydney Informatics Hub at the University of Sydney.

## Author Contributions

K. R., J. H., K.V.B. and E.C.H. conceptualized the study.; E. M., J. M., J. R., M. P., E. M. M. and K.V.B. performed the research; E. M., L.L., J. C. O. M., M. P. and K.V.B. analysed the data; E.C.H. provided supervision; E. M., K.V.B. and E.C.H. wrote the original draft. All authors reviewed and approved the final manuscript.

## Competing Interest Statement

The authors declare no completing interests.

## Notes

### Competing Interest Statement

The authors have declared no competing interest.

