## Supplementary Information for "Hidden diversity and expanded host range of sarthroviruses, including terrestrial vertebrates"

**Supplementary Material**

**Table S1.** Sample, location, sampling date and host information for each library analysed in this study.

| # of individuals | Sample type | Host source | Library | Location | Date |
|---|---|---|---|---|---|
| 16581.9 | Liver | *H. garnotii* | Gecko 1.1 | Freshwater, Sydney | 2025 |
| | Lung | | Gecko 1.2 | | |
| 16558.3<br>16558.4 | Liver | | Gecko 2.1 | Cremorne, Sydney | 2025 |
| 16558.1<br>16558.2<br>16558.3<br>16558.4 | Lung | | Gecko 2.2 | | |
| 16558.8 | Liver | | Not sequenced | Manly, Sydney | 2026 |
| 14364.1 | Faeces | *A. pusillus doriferus* | Seal 1 | Avoca beach, Central Coast | 2021 |
| SS001 | Whole body | Demospongiae | Sponge 1 | Chowder Bay, Sydney | 2023 |
| SS002<br>SS003 | Whole body | Demospongiae | Sponge 2 | Chowder Bay, Sydney | 2023 |
| SS004 | Whole body | Demospongiae | Sponge 3 | Chowder Bay, Sydney | 2023 |
| NA | Liver and gill | *Cyprinus carpio* | SRR13709132* (10) | Coomealla, Murray River | 2022 |
| NA | Faeces | *Canis lupus dingo* | SRR32313011* (46) | Myall Lake, Mid North Coast | 2022 |
| NA | Marine sediment | Unknown | SRR8820974* | Bare Island, Sydney | 2016 |
| NA | Marine sediment | Unknown | SRR8821107* | Bare Island, Sydney | 2016 |
| NA | Marine sediment | Unknown | SRR8820979* | Balls Head, Sydney | 2016 |
| NA | Marine sediment | Unknown | SRR8820972* | Bare Island, Sydney | 2016 |

| | | | | | |
|---|---|---|---|---|---|
| NA | Marine sediment | Unknown | SRR8820893* | Towra Point, Sydney | 2016 |
| NA | Marine sediment | Unknown | SRR8820786* | Cobblers Beach, Sydney | 2016 |
| NA | Marine sediment | Unknown | SRR8820861* | Balls Head, Sydney | 2016 |
| NA | Marine sediment | Unknown | SRR8820785* | Cobblers Beach, Sydney | 2016 |
| NA | Marine sediment | Unknown | SRR8820899* | Towra Point, Sydney | 2016 |
| NA | Marine sediment | Unknown | SRR8820976* | Balls Head, Sydney | 2016 |
| NA | Marine sediment | Unknown | SRR6401741* | Balls Head, Sydney | 2016 |
| NA | Marine sediment | Unknown | SRR8820897* | Salmon Haul, Sydney | 2016 |
| NA | Marine sediment | Unknown | SRR8820831* | Cobblers Beach, Sydney | 2016 |
| NA | Marine sediment | Unknown | SRR8820788* | Cobblers Beach, Sydney | 2016 |
| NA | Marine sediment | Unknown | SRR6401740* | Bare Island, Sydney | 2016 |
| NA | Marine sediment | Unknown | SRR8820918* | Lilli Pilli Pool, Sydney | 2016 |
| NA | Marine sediment | Unknown | SRR8821045* | Balls Head, Sydney | 2016 |
| NA | Marine sediment | Unknown | SRR8820973* | Bare Island, Sydney | 2016 |
| NA | Marine sediment | Unknown | SRR8820970* | Bare Island, Sydney | 2016 |

*Libraries downloaded from the NCBI SRA database

**Table S2**. Primers used to screen and complete partial viruses

| **Virus** | **Forward primer (5' - 3')** | **Reverse primer (5' - 3')** |
|---|---|---|
| Itsy bitsy virus | Sarthro-F<br>TATCCGGTTAAAATGTCTTGTC | Sarthro-R<br>AACTATGTTGTACACGGTGAC |
| Seal faecal associated nodavirus 2 | Seal-noda-F1<br>CTGTTACCATGTTCACCGTT<br>Seal-noda-F2<br>CCCAAATTACCCTAGGAACAT<br>T<br>Seal-noda-F3<br>GAGTATACCTCGATCCTTGG | Seal-noda-R1<br>ATCTTAGGTATTATGTGCTCGC<br>Seal-noda-R2<br>AAGGTATGAGACGAAAGCATT<br>Seal-noda-R3<br>CTACAACGTCATTCCAGTCAT |
| Indo-pacific nodavirus 2 | Gecko-noda-F1<br>CGCCTGTTGATGAGAAGAAC<br>Gecko-noda-F2<br>ACACAGTTGTCACATTGGA | Gecko-noda-F1<br>CAGATTTTGGATGCGAATGTAT<br>Gecko-noda-F2<br>GTAACGCACAGTACACATTAA<br>A |
| Gecko dicistroviru<br>s | Gecko-dicistro-F<br>CGCTACGTGTAAGAATTGTAA<br>T | Gecko-dicistro-R<br>ATCGCGTTTCCATTAAAATCAT |

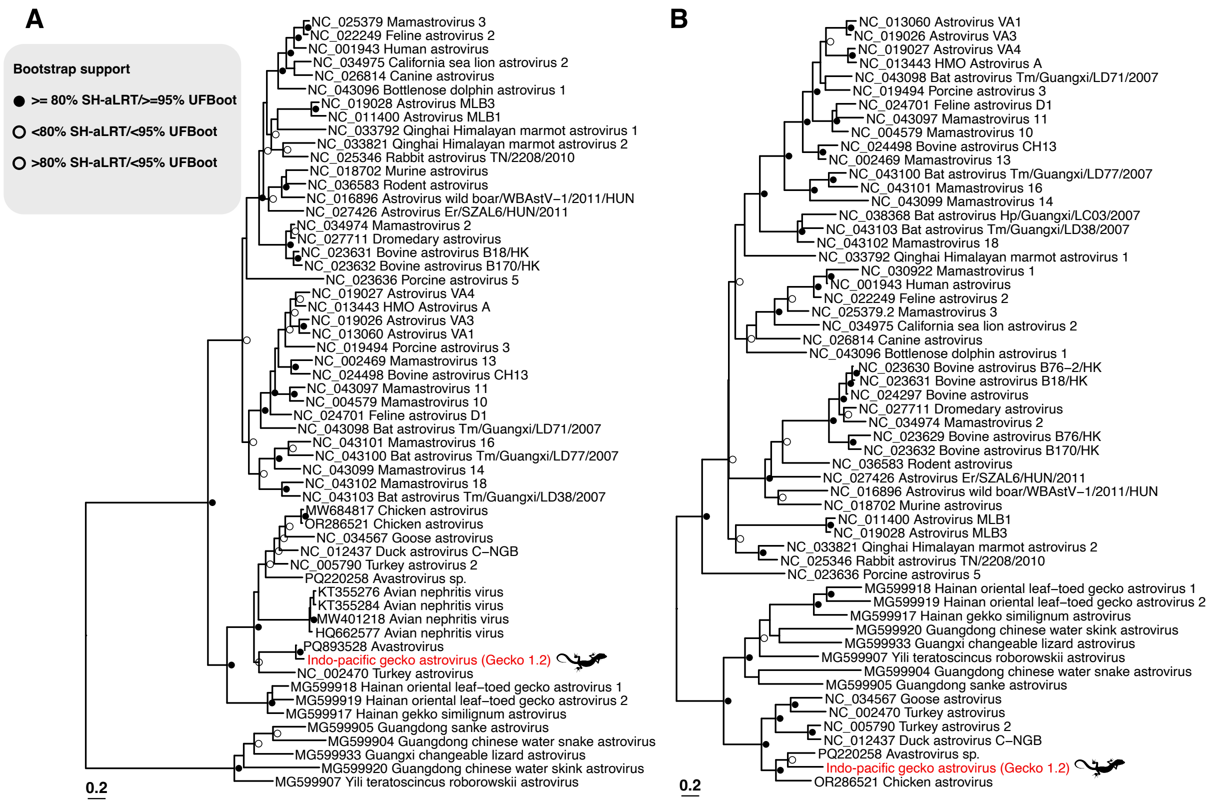


**Figure S1. Phylogenetic relationships among the *Astroviridae*.** Maximum likelihood phylogenetic trees of the (A) RdRp and (B) capsid MAFFT alignment of available NCBI GenBank astrovirus sequences and the gecko astrovirus sequences identified in this study (shown in red). Unaligned sections and gaps were removed from the amino acid alignments before tree estimation. All horizontal branch lengths are drawn to a scale of substitutions per site and the support values (SH-aLRT and UFBoot) are shown at the nodes. The tree is mid-pointed for clarity.

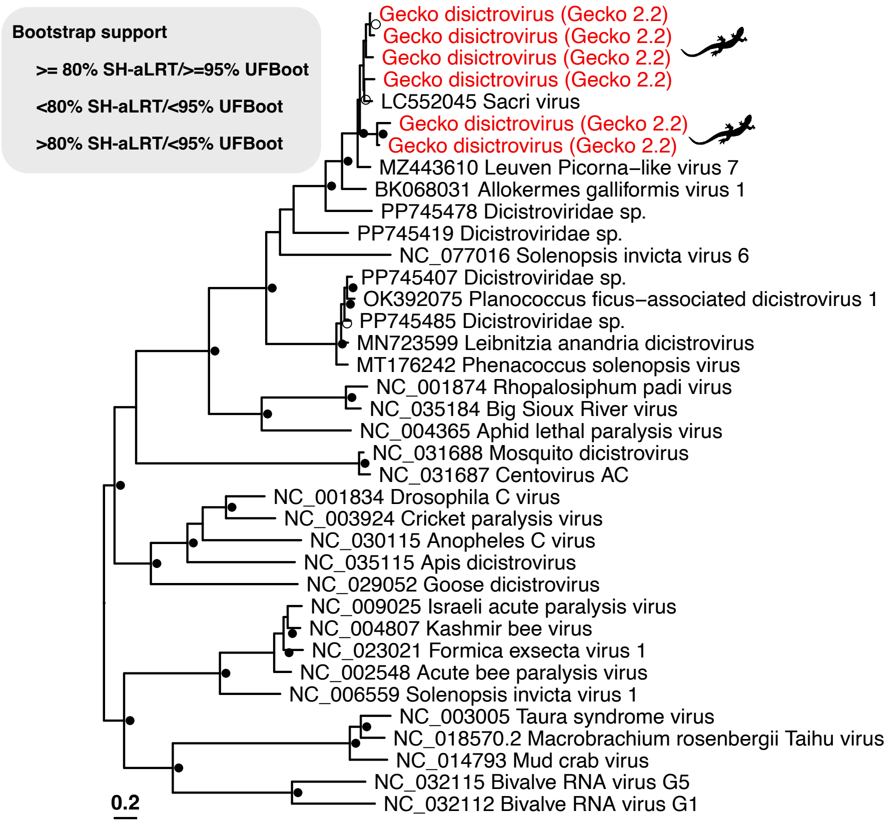


**Figure S2. Phylogenetic relationships among the *Dicistroviridae.*** Maximum likelihood phylogenetic tree of the non-structural gene MAFFT alignment of available NCBI GenBank astrovirus sequences and the gecko astroviruses identified in this study (shown in red). Unaligned sections and gaps were removed from the amino acid alignments before tree estimation. All horizontal branch lengths are drawn to a scale of substitutions per site and the support values (SH-aLRT and UFBoot) are shown at the nodes. The tree is mid-pointed for clarity.

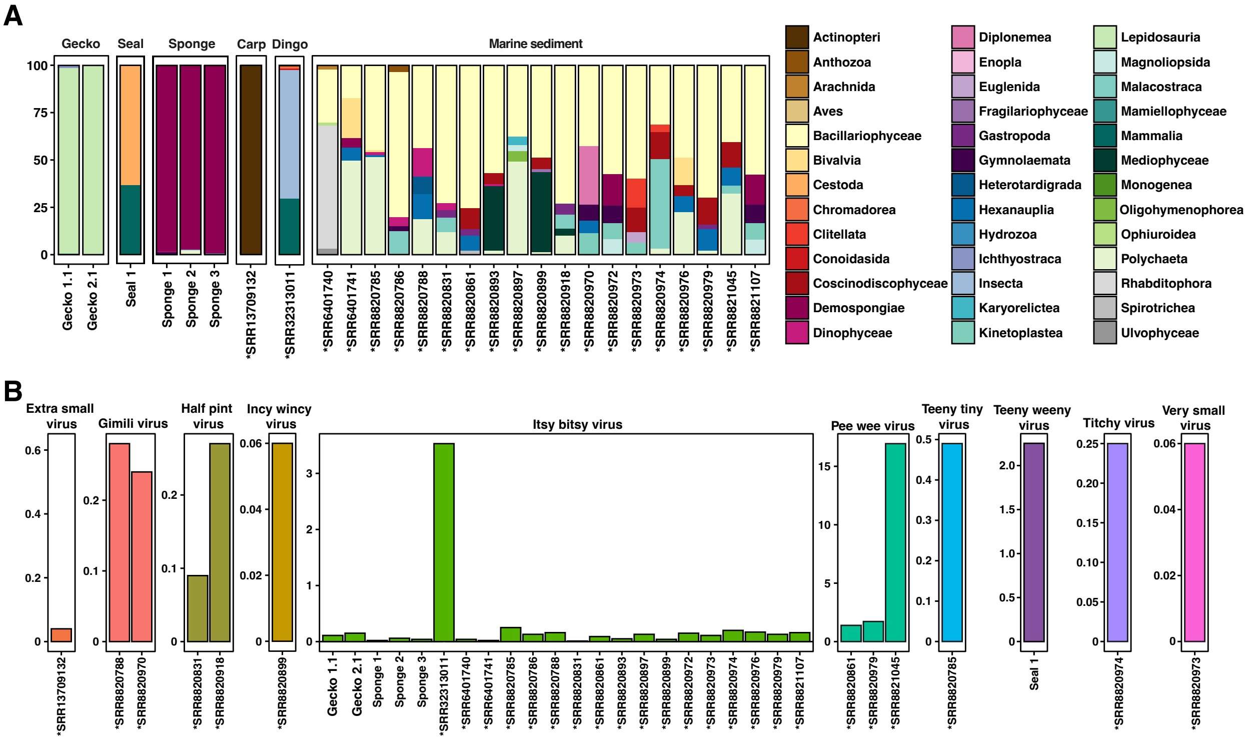


**Figure S3. Taxonomic classification using Kraken2 and virus abundance.** (A) Eukaryota reads per library as classified by Kraken2. For better resolution the graph only represents the top 5 eukaryote hits for each library. (B) Read abundances of each sarthrovirus is given as the number of reads per million (RPM) and calculated using RSEM. The libraries downloaded from the NCBI SRA database are denoted by an asterisk.
